# A Precision Medicine Approach to Metabolic Therapy for Breast Cancer in Mice

**DOI:** 10.1101/2021.12.15.472809

**Authors:** Ngozi D. Akingbesote, Aaron Norman, Wanling Zhu, Alexandra A. Halberstam, Xinyi Zhang, Julia R. Foldi, Maryam B. Lustberg, Rachel J. Perry

## Abstract

Increasing evidence highlights the possibility for approaches targeting metabolism as potential adjuvants to cancer therapy. Sodium-glucose transport protein 2 (SGLT2) inhibitors are the newest class of antihyperglycemic therapies, and have recently been highlighted as a novel therapeutic approach to breast cancer. To our knowledge, however, SGLT2 inhibitors have not been applied in the neoadjuvant setting as a precision medicine approach to combining metabolic therapy with standard of care therapy for this devastating disease. In this study we combine the SGLT2 inhibitor dapagliflozin with paclitaxel chemotherapy in both lean and obese mice. We show that dapagliflozin enhances the efficacy of paclitaxel, reducing tumor glucose uptake and prolonging survival in an insulin-dependent manner in some but not all breast tumors. Our data find a genetic signature for breast tumors most likely to respond to dapagliflozin in combination with paclitaxel. Tumors driven by mutations upstream of canonical insulin signaling pathways are likely to respond to such treatment, whereas tumors driven by mutations downstream of canonical insulin signaling are not. These data demonstrate that dapagliflozin enhances the response to chemotherapy in mice with breast cancer and suggest that breast cancer patients with driver mutations upstream of canonical insulin signaling may be most likely to benefit from this neoadjuvant approach. A clinical trial is currently in preparation, with an application recently submitted for Yale Human Investigations Committee approval, to test this hypothesis in breast cancer patients.

**One Sentence Summary:** We identify a driver mutation signature by which glucose-wasting metabolic therapy (dapagliflozin) enhances the efficacy of chemotherapy in mice with breast cancer.

## INTRODUCTION

Breast cancer is the second leading cause of cancer deaths in women, with one in eight women in the U.S. developing this disease over her lifetime. The incidence of breast cancer is projected to increase more than 50% between 2011 and 2030 *(1)*. This troubling prediction is attributable in part to rising rates of obesity, which accelerates the appearance and progression of breast cancer in postmenopausal women *(2, 3)*. Similarly, primary tumor appearance *(4–6)*, volume *(5, 7–9)*, and metastasis *(7–11)* increase with a high-calorie diet in implanted, genetic, and chemically-induced mouse models of breast cancer.

Metabolism-targeting neoadjuvant approaches have gained increasing popularity in breast cancer treatment. The biguanide metformin, an inhibitor of gluconeogenesis *(12–14)* and the most prescribed antidiabetes drug worldwide *(15)*, has been by far the most popular antihyperglycemic approach used for cancer. At supra-pharmacologic doses – several orders of magnitude higher than in human patients – metformin is capable of reducing breast cancer cell proliferation *(16–19)*, and slowing tumor growth *in vivo* in rodents *(20–22)*. Metformin is safe and shows good target engagement in women with breast cancer. Studies regarding its efficacy, however, have been mixed, with most *(23–29)*, but not all, adjuvant and neoadjuvant studies *(30–32)* failing to observe differences in tumor cell proliferation markers or in outcomes in human patients.

Preclinical data highlight the newest class of diabetes drug, SGLT2 inhibitors, as a potentially attractive alternative to metformin therapy for breast cancer. As metformin acts primarily to inhibit gluconeogenesis, its ability to lower blood glucose and insulin concentrations is likely minimal under postprandial conditions in individuals without diabetes. SGLT2 inhibitors, by contrast, cause glycosuria and, therefore, glucose wasting under both fasting and postprandial conditions. We recently demonstrated that the SGLT2 inhibitor dapagliflozin slows breast tumor growth when administered as monotherapy in obese mice, and that its anticancer effect was reliant upon its ability to reverse fasting hyperinsulinemia *(33)*. A direct effect of SGLT2 inhibition in tumors cannot be ruled out, because data from the Human Protein Atlas (https://www.proteinatlas.org/about/licence) indicate that breast tumors do express SGLT2, albeit at low levels *(34)*. However, considering data demonstrating that chronically infusing insulin to abrogate the insulin-lowering effects of dapagliflozin fully abrogated its beneficial effects, it is most likely that the tumor suppressive effects of dapagliflozin are likely attributable to its ability to reduce insulin-stimulated tumor glucose uptake *(33)*. However, to our knowledge, SGLT2 inhibitors have never been administered in combination with chemotherapy in *in vivo* breast cancer models. This is a point of substantial clinical importance because breast cancer treatment regimens are effective, but not unanimously so: response rates are particularly low in estrogen receptor-positive breast cancer *(35)*. Thus, new neoadjuvant approaches are critical to improve the efficacy of standard-of-care breast cancer treatment.

In addition, at a time when precision medicine approaches are gaining increasing popularity in selecting targeted therapies for patients. Such approaches, however, have largely failed to investigate and target metabolism. In this study, we aimed to generate a genetic signature for breast tumors responsive to dapagliflozin as an adjunct to chemotherapy. *In vivo* studies in seven murine models of breast cancer with different driver mutations demonstrated a genetic signature of those tumors that responded to dapagliflozin: in tumors with driver mutations upstream of the insulin receptor, dapagliflozin improved the efficacy of paclitaxel, whereas in tumors with driver mutations downstream of the insulin receptor, dapagliflozin was ineffective. These data predict that tumor genetics may be utilized to design metabolism-targeting neoadjuvant treatments for patients with hyperinsulinemia.

## RESULTS

### Dapagliflozin reduces insulin more than other common diabetes drugs

Our previous studies have shown plasma insulin concentrations to be a predictor of tumor growth rates in colon *(36)* and breast cancer *(33)*. Therefore, we first aimed to determine which commonly prescribed antihyperglycemic medications would most significantly reduce both fasting and postprandial plasma insulin concentrations. The biguanide metformin, the glucagon-like peptide-1 agonist semaglutide, the thiazolidinedione pioglitazone, and the sulfonylurea glipizide all showed some ability to lower plasma glucose concentrations. The SGLT2 inhibitor dapagliflozin, however, lowered plasma insulin more substantially than any of the other agents in healthy mice. Moreover, dapagliflozin was the only agent of the five tested to lower both fasting and postprandial plasma insulin (Figure 1A-B). For this reason, we selected dapagliflozin for future studies in murine models of breast cancer.

**Fig. 1.**
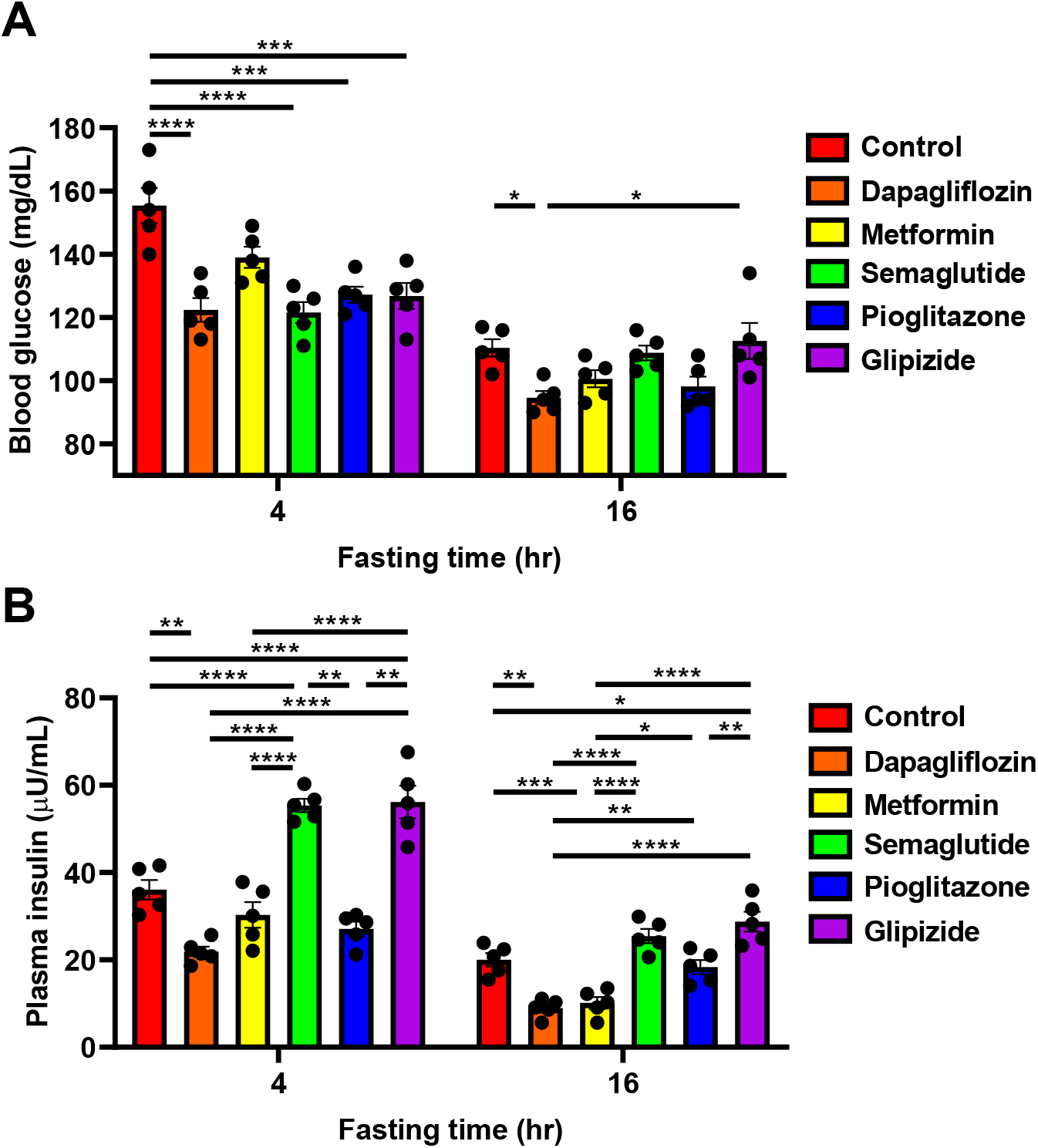
The SGLT2 inhibitor dapagliflozin outperforms diabetes drugs in other classes at lowering both fasting and postprandial plasma insulin concentrations. (A) Blood glucose. (B) Plasma insulin. In both panels, \**P*<0.05, \*\**P*<0.01, \*\*\**P*<0.001, \*\*\*\**P*<0.0001 by ANOVA with Tukey’s multiple comparisons test. n=5 per group.

### Dapagliflozin slows tumor growth in lean and obese MMTV-PyMT mice

In previous work, we have shown that dapagliflozin slowed E0771 breast cancer growth when applied as monotherapy in obese mice (29). Here, we examined for the first time whether dapagliflozin would slow tumor growth in lean MMTV-PyMT mice, a murine breast cancer model driven by the polyoma virus middle T antigen and commonly expressing an Erbb2 mutation. We found that, as expected, SGLT2 inhibition caused glucose wasting in urine, suppressing weight gain without causing ketosis (Supplementary Figure 1A-C). This glucose wasting resulted in both chronic and acute (within 48 hours, before mice diverged in body weight) reductions in plasma glucose and insulin concentrations in recently fed (6 hr fasted) mice given ad lib access to both regular chow and a high-fat, high-carbohydrate Western diet. These reductions in plasma insulin concentrations inhibited tumor glucose uptake: both acute and chronic dapagliflozin treatment lowered MMTV-PyMT tumor [14C] 2-deoxyglucose uptake, with chronic treatment more markedly reducing glucose uptake than acute (Figure 2A-E, Supplementary Figure 1D). These reductions in tumor glucose uptake correlated with a profound effect of metabolic manipulation on tumor growth: whereas both glucose uptake and tumor growth were accelerated in Western diet fed mice as compared to chow fed, dapagliflozin slowed tumor growth in both lean and obese animals (Figure 2F).

**Fig. 2.**
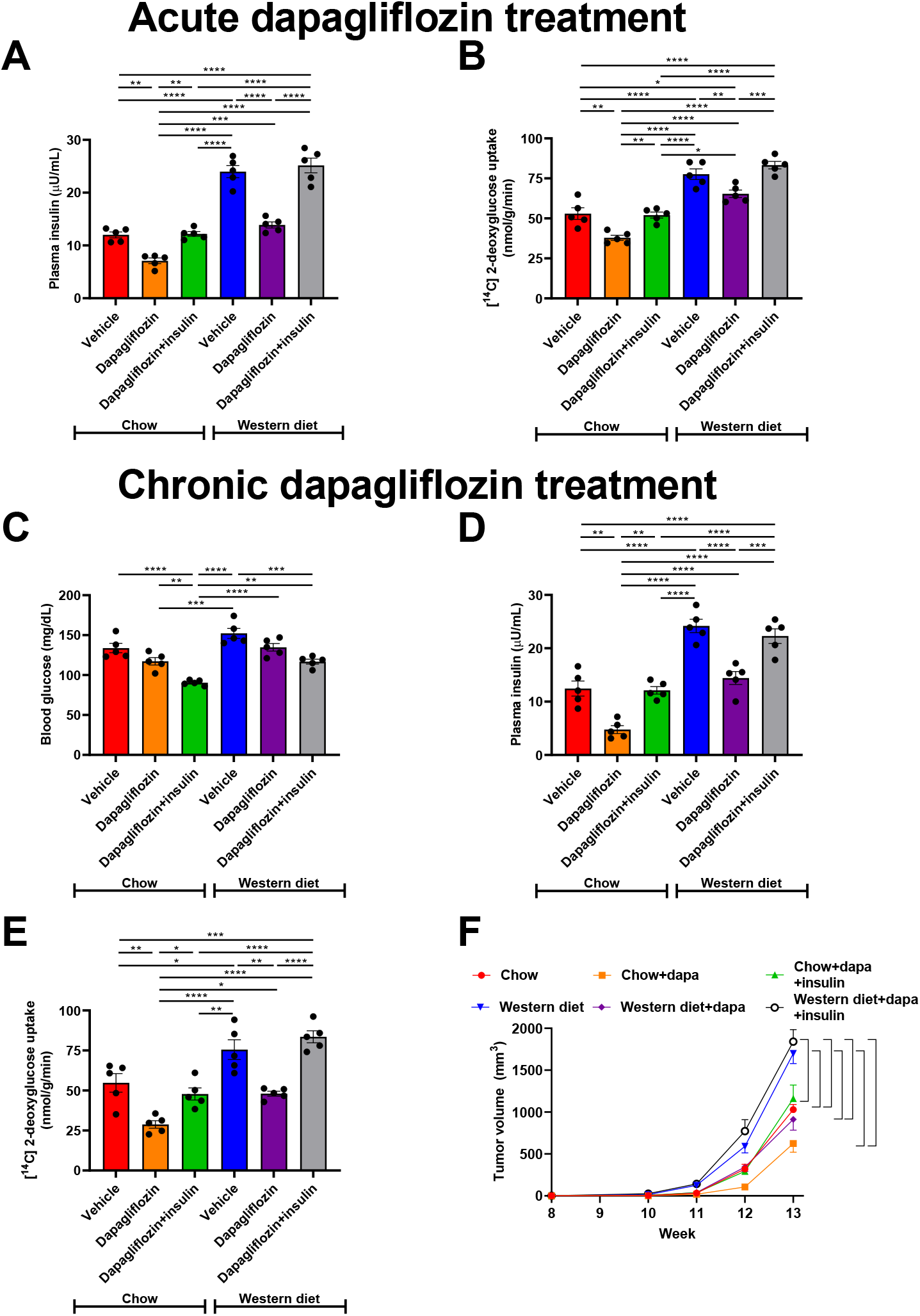
Dapagliflozin slows spontaneous tumor growth in MMTV-PyMT mice through its effect to lower plasma insulin in lean and obese mice. (A) Plasma insulin and (B) Tumor [^14^C] 2-deoxyglucose uptake in mice treated acutely with a single dose of dapagliflozin. (C)-(D) Blood glucose and plasma insulin in mice treated continuously for four weeks with dapagliflozin. (D)-(E) Tumor [^14^C] 2-deoxyglucose uptake in mice treated chronically with dapagliflozin. In panels (A)-(E), \**P*<0.05, \*\**P*<0.01, \*\*\**P*<0.001, \*\*\*\**P*<0.0001 by ANOVA with Tukey’s multiple comparisons test. (F) Tumor growth. Data are the mean±S.E.M. of n=5 per group. The brackets denote statistically significant (P<0.05) comparisons by ANOVA with Tukey’s multiple comparisons test.

### Dapagliflozin slows 4T1 tumor growth in lean and obese mice

Next, we studied a second murine tumor model, 4T1 subcutaneous breast cancer, which is driven by a p53 mutation (34,35). BALB/c mice treated with Western diet did become overweight (Supplementary Figure 2A), and both hyperglycemic and hyperinsulinemic but not ketotic (Figure 3A-B, Supplementary Figure 2B). Western diet enhanced both glucose uptake and tumor growth, but dapagliflozin reduced each of these parameters in both chow and Western diet fed mice (Figure 3C-D). Employing chronic insulin infusion to prevent the effect of dapagliflozin to reduce plasma insulin concentrations, however, abrogated its effect to slow tumor growth in both lean and obese mice. This demonstrated that dapagliflozin primarily slows 4T1 tumor growth by lowering circulating insulin concentrations. Reduced adiposity in dapagliflozin-treated mice is unlikely to explain slower MMTV-PyMT tumor growth, as incubating 4T1 cells in dapagliflozin increased cell number rather than decreasing it (Supplementary Figure 2C). Supplying physiologic concentrations of saturated fatty acid (palmitate) slowed cell division. Inhibiting fatty acid metabolism by coculturing with etomoxir partially rescued the effect of palmitate to slow cell division (Supplementary Figure 2D). Taken together, these data argue against a direct effect of dapagliflozin to inhibit, or adipose-derived fatty acids to promote, breast cancer growth.

**Fig. 3.**
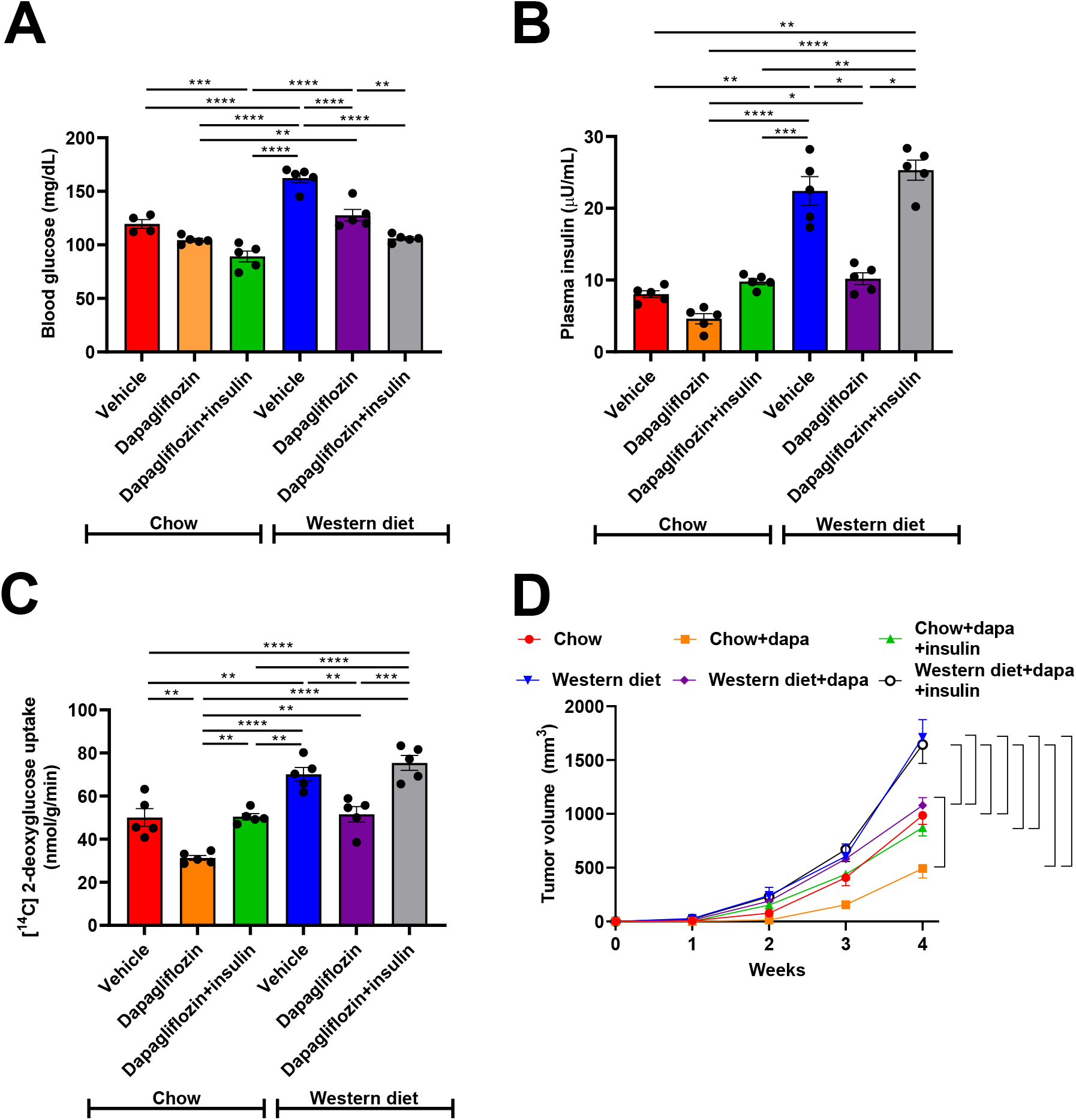
Dapagliflozin slows orthotopic 4T1 tumor growth by lowering plasma insulin in lean and obese mice. (A) Blood glucose. (B) Plasma insulin. (C) Tumor [14C] 2-deoxyglucose uptake. In panels (A)-(C), \**P*<0.05, \*\**P*<0.01, \*\*\**P*<0.001, \*\*\*\**P*<0.0001 by ANOVA with Tukey’s multiple comparisons test. (D) Tumor growth. Data are the mean±S.E.M. of n=5 per group. The brackets denote statistically significant (*P*<0.05) comparisons by ANOVA with Tukey’s multiple comparisons test.

### Dapagliflozin enhances efficacy of chemotherapy in mice with breast cancer

Although our and others’ preclinical data demonstrating an effect of SGLT2 inhibitors to slow tumor growth are promising, dapagliflozin should and will never be administered as monotherapy for breast cancer. We next examined the impact of dapagliflozin in mice treated with standard-of-care paclitaxel. Both paclitaxel and dapagliflozin reduced tumor glucose uptake and improved survival in both MMTV-PyMT and 4T1 breast cancer models. However, dapagliflozin and paclitaxel showed synergistic effects against breast cancer: the two drugs showed additive effects to reduce tumor glucose uptake and improve survival in MMTV-PyMT and 4T1 tumor-bearing mice (Figure 4A-D).

**Fig. 4.**
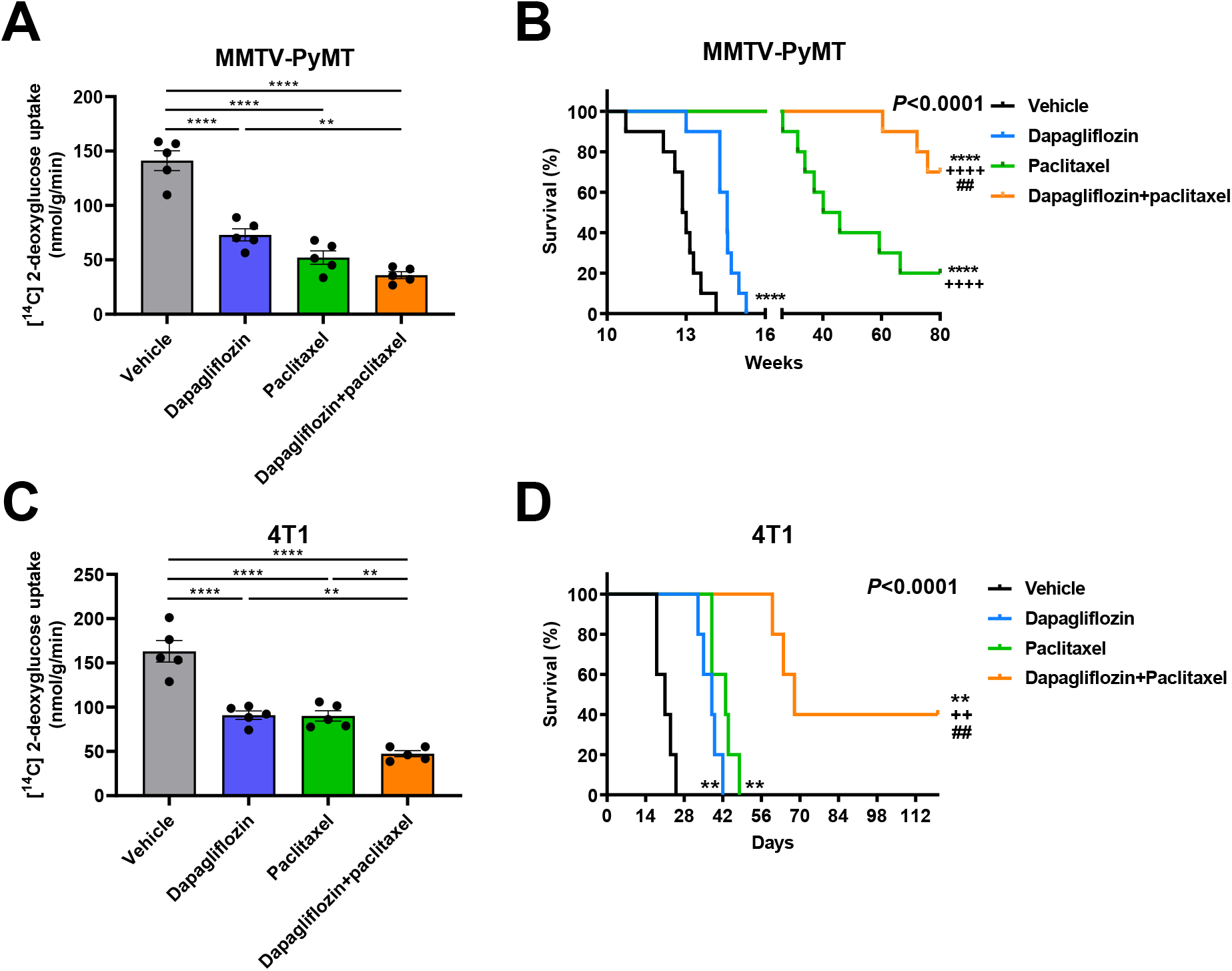
Dapagliflozin enhances the efficacy of chemotherapy in two murine models of breast cancer. (A) Tumor glucose uptake and (B) Survival in MMTV-PyMT tumor-bearing mice. The x-axis refers to weeks of life. (C) Tumor glucose uptake and (D) Survival in 4T1 tumor-bearing mice. The x-axis refers to days after tumor cell injection. In panels (A) and (C), \*\**P*<0.01, \*\*\*\**P*<0.0001. In panels (B) and (D), \*\**P*<0.01, \*\*\*\**P*<0.0001 vs. vehicle, ^++^*P*<0.01, ^++++^*P*<0.0001 vs. dapagliflozin, ##P<0.01 vs. paclitaxel via the Mantel-Cox log-rank test, adjusted for multiple comparisons. The p-values in the upper right corner of each survival curve refer to the overall curve comparison using the Mantel-Cox log-rank test.

### Dapagliflozin does not cause fatigue, weight loss, or neuropathy

In considering dapagliflozin as a potential neoadjuvant for future study in humans, it is crucial to examine the possibility that it may enhance existing side effects of paclitaxel chemotherapy. To that end, we assessed the impact of dapagliflozin on anorexia and on neuropathy, both common adverse effects of chemotherapy treatment. Encouragingly, we found that dapagliflozin did not cause fatigue as indicated by spontaneous activity, reduce food intake, or alter thermal latency as monotherapy or when added to chemotherapy in either MMTV-PyMT or 4T1 tumor-bearing mice (Figure 5A-H). This demonstrates that dapagliflozin might hold promise as a safe and effective adjuvant to standard-of-care chemotherapy in breast cancer.

**Fig. 5.**
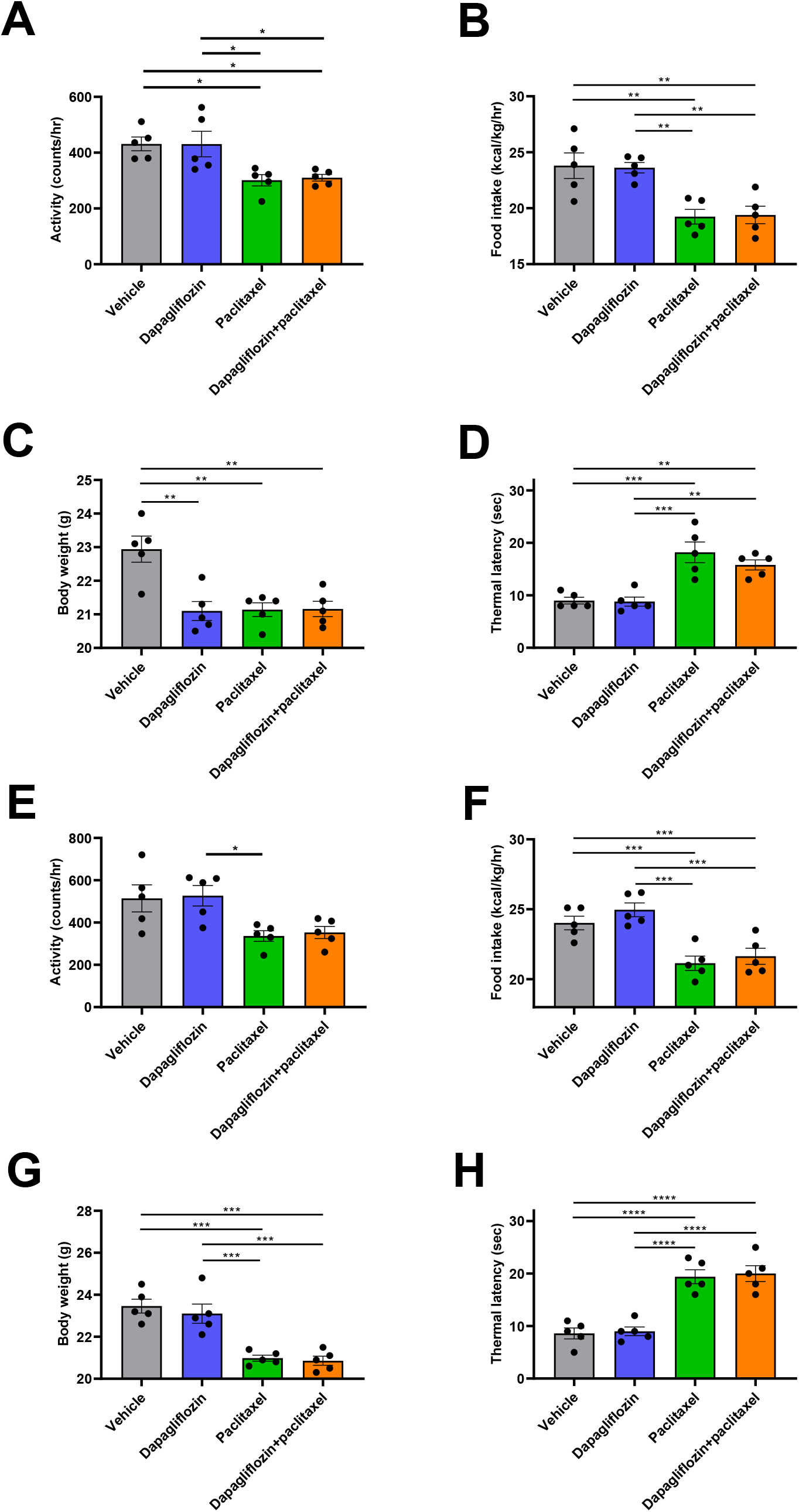
Dapagliflozin has no adverse effects to cause fatigue, anorexia, weight loss, or neuropathy in MMTV-PyMT or 4T1 tumor-bearing mice. (A)-(C) Spontaneous activity, food intake, and thermal latency during week 3 of chemotherapy or vehicle treatment. (D) Body weight after 4 weeks in MMTV-PyMT mice. (E)-(G) Spontaneous activity, food intake, and thermal latency during week 3 of chemotherapy or vehicle treatment. (H) Body weight after 4 weeks in 4T1 tumor-bearing mice. In all panels, \**P*<0.05, \*\**P*<0.01, \*\*\**P*<0.001 by ANOVA with Tukey’s multiple comparisons test.

### Driver mutations predict the response to dapagliflozin in breast cancer

Precision medicine is an increasingly attractive arena. To determine whether certain tumor drivers may be more likely than others to respond to dapagliflozin, we examined the impact of this agent on survival in obese mice with tumors driven by five additional mutation profiles (Supplementary Table 2). We observed a genetic signature of tumors responding to SGLT2 inhibition. While dapagliflozin did not synergize with paclitaxel in mice with p53-driven M6, MYC-driven M158 tumors, or MEK1-driven EpH4 1424 tumors with Brca1-driven tumors, it prolonged survival in mice with Pten-driven EMT6 tumors and HRAS-driven Ac711 tumors (Figure 6A-E), in addition to ERBB2-driven MMTV-PyMT, and p53 and Pik3ca-driven 4T1 tumors (Figure 4B, Figure 4D). Taken together, these data suggest that tumors with mutations upstream of canonical insulin signaling responded to dapagliflozin as an adjuvant to chemotherapy, whereas tumors driven by mutations downstream of canonical insulin signaling did not (Figure 7). These data highlight opportunities for the development of precision medicine approaches to utilize metabolic adjunct therapies for breast cancer informed by tumor genetics.

**Fig. 6.**
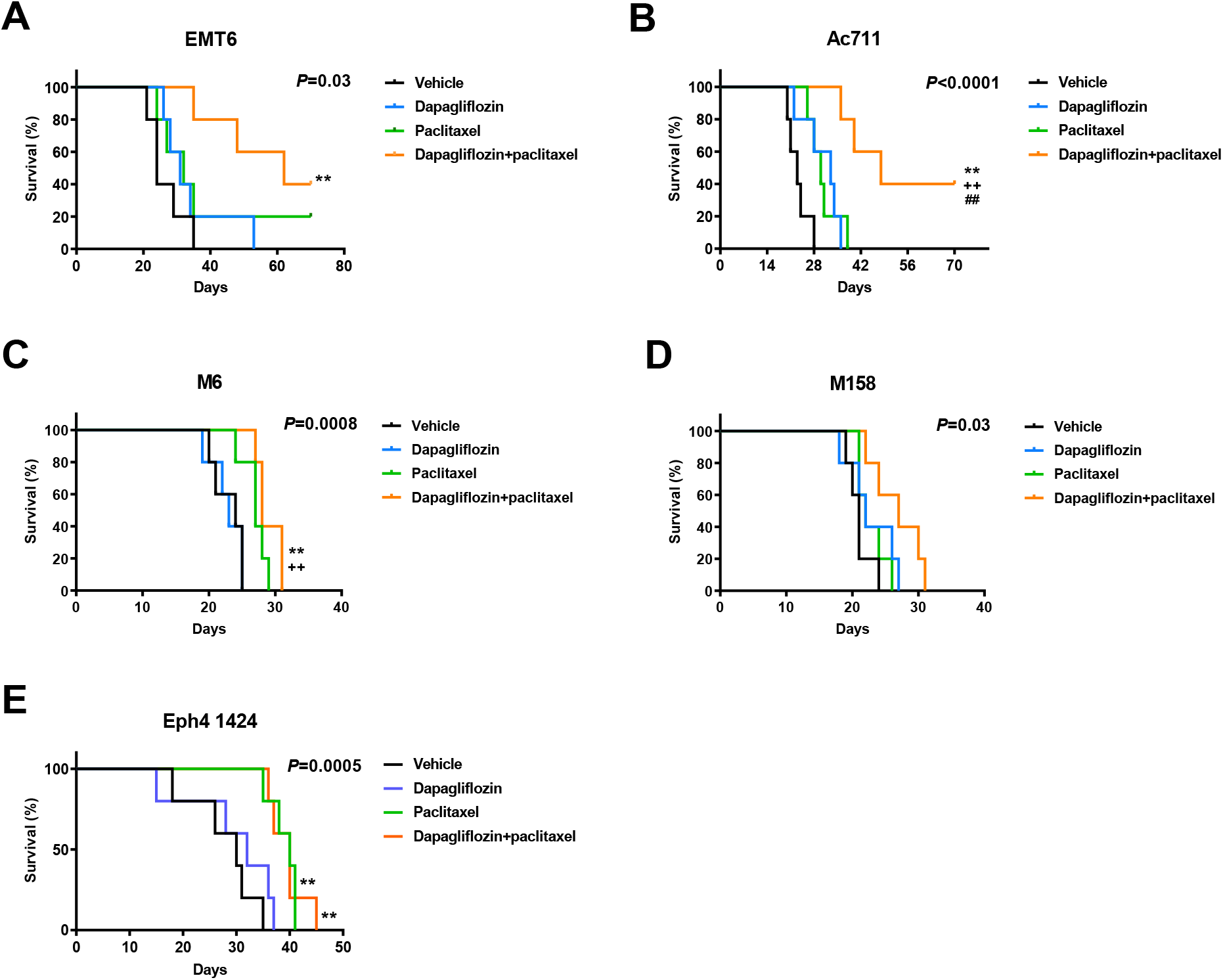
Tumor drivers predict the response to dapagliflozin as an adjunct to paclitaxel. (A)-(E) Survival in EMT6, Ac711, M6, M158, and Eph 1424 tumor-bearing mice, respectively. \*\**P*<0.01 vs. vehicle, ++*P*<0.01 vs. dapagliflozin, ##*P*<0.01 vs. paclitaxel by the Mantel-Cox log-rank test, adjusted for multiple comparisons. The p-values in the upper right corner of each survival curve refer to the overall curve comparison using the Mantel-Cox log-rank test.

**Fig. 7.**
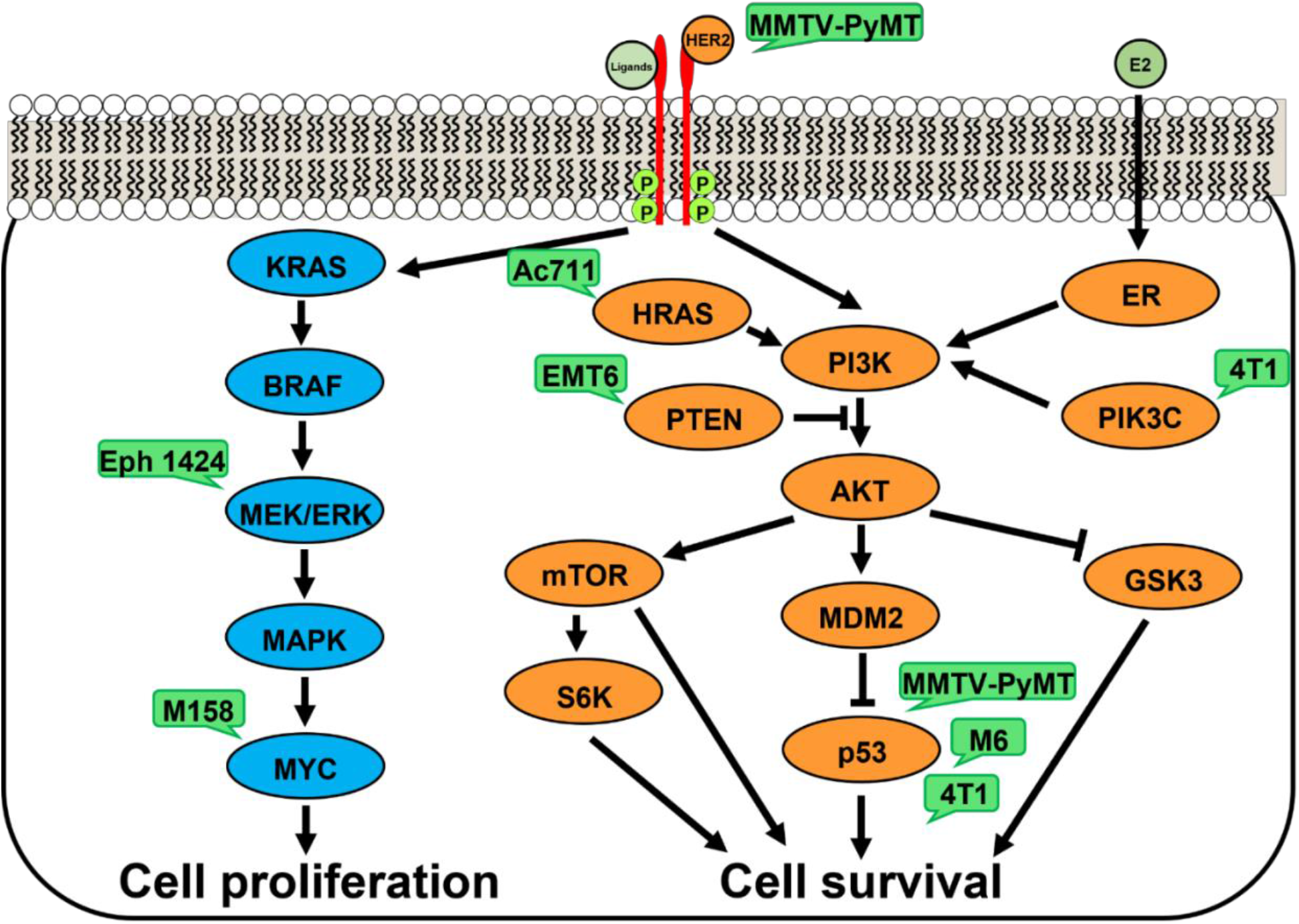
Driver mutations tested in this study. Green bubbles show models utilized herein.

## DISCUSSION

Despite the undisputed epidemiologic link between excess weight and numerous cancers, including breast cancer, clinical trials examining the effect of metabolic therapy (almost exclusively, metformin) against cancer have been under-pursued and underwhelming. This may be at least in part to metformin’s underwhelming effect to reduce plasma insulin concentrations by less than 20% *(37, 38)* (Figure 1B). Given its higher potential efficacy, in this study, we examined the impact of dapagliflozin as an adjunct in combination with chemotherapy. We treated mice with breast cancer driven by a complement of mutations with dapagliflozin in combination with paclitaxel chemotherapy.

Importantly, without enhancing adverse effects of paclitaxel, dapagliflozin improved the efficacy of paclitaxel to promote neuropathy, but it improved the efficacy of chemotherapy to slow tumor growth in murine breast cancer models with mutations in pathways upstream, but not downstream, of canonical insulin signaling (Figure 7). These data suggest that precision medicine approaches to neoadjuvant treatment are of pressing importance. Indeed, a recent meta-analysis hinted at a differing effect of obesity on outcomes based on cancer subtype: Lohmann and colleagues observed a larger magnitude of the deleterious effect of excess body weight in triple-negative and hormone receptor-positive, HER2-negative breast cancer as compared to hormone receptor-positive, HER2-positive breast cancer *(39)*. However, these data stop short of suggesting specific breast cancer driver mutations that may be more or less responsive to therapies aiming to reduce circulating insulin concentrations. Therefore, in this study we aimed to determine a genetic signature for responsiveness to dapagliflozin in mice with breast cancer.

The primary advance of the current study is its comprehensive assessment of which genetic drivers of breast cancer may be most responsive to SGLT2 inhibitors and perhaps other insulin-lowering agents. Our data demonstrate a genetic signature for tumors in which dapagliflozin improved the response to chemotherapy: E0771, 4T1, EMT6, and Ac711 tumors are driven by proteins upstream of PI3K/Akt, key mediators of insulin signaling in health *(40)* and disease *(41, 42)*. Dapagliflozin enhanced the efficacy of paclitaxel in these tumors. By contrast, SGLT2 inhibition did not improve chemotherapy efficacy in tumors driven by mutations in the KRAS/MEK/ERK cell proliferation pathway (M158 and EpH4 1424), or p53 mutations downstream of PI3K/Akt in the cell survival pathway (M6).

Further emphasizing the physiologic relevance of dapagliflozin’s inhibition of the insulin signaling pathway, matching plasma insulin concentrations in dapagliflozin-treated 4T1 and MMTV-PyMT tumor-bearing mice to concentrations measured in untreated controls abrogated the beneficial effect of dapagliflozin to enhance the efficacy of chemotherapy in these animals. While we cannot completely rule out a direct effect of dapagliflozin to reduce tumor glucose uptake independently of insulin, two data sets argue against this interpretation. First, we observed no difference in tumor growth rates between untreated controls and mice treated with dapagliflozin plus insulin (Figures 2F, 3D). This suggests that adding insulin undermines the efficacy of the SGLT2 inhibitor in slowing tumor growth. If the SGLT2 inhibitor were acting directly to reduce tumor glucose uptake, insulin infusion would not be expected to fully abrogate the effect of dapagliflozin. Second, the mutational signature of tumors responsive to dapagliflozin – that is, those with mutations downstream of canonical insulin signaling – argues against a direct effect of dapagliflozin on the tumor. If the “direct action” hypothesis were correct, one would predict a similar effect of SGLT2 inhibition to slow tumor growth regardless of whether the driver mutation is related to insulin and/or glucose metabolism.

An additional key point in this study is the efficacy of dapagliflozin in lean animals. Although both canagliflozin *(33, 43, 44)*, dapagliflozin *(33)*, empagliflozin *(45)*, and impragliflozin *(46)* have been shown to reduce breast cancer cell division *in vitro* under high glucose conditions, the efficacy of any SGLT2 inhibitor has rarely been examined *in vitro* in glucose and insulin concentrations within the physiologic range. Even fewer studies have been performed to examine the efficacy of SGLT2 inhibitors *in vivo* in lean animals. A single study reported use of dapagliflozin as anti-breast cancer therapy in regular chow fed, athymic, nude mice with MCF-7 breast cancer xenografts *(44)*. While dapagliflozin was effective in slowing tumor growth in this model, the absence of a typical anticancer immune response in this model leaves questions as to the potential efficacy of this agent *in vivo* in those with intact immune systems and without obesity.

Relatedly, fasting hyperinsulinemia can occur in those without diabetes and even without impaired glucose tolerance *(47)* and is rapidly on the rise in the U.S. and worldwide. Between the National Health and Nutrition Examination Surveys conducted between 1988–1994 and 1999–2002, for instance, the age-adjusted incidence of fasting hyperinsulinemia increased by 35% *(48)*. In the last two decades, the prevalence of hyperinsulinemia even in those without diabetes has increased, while the prevalence of diabetes also increased by close to 50% in the same time period *(49)*. These data emphasize that there is a critical need to understand therapies that might work for both those with diabetes and those with hyperinsulinemia without diabetes.

In summary, we demonstrate here that the SGLT2 inhibitor dapagliflozin improves the efficacy of chemotherapy to slow breast tumor growth in a tumor driver-dependent manner. We reveal that mice with breast cancer driven by mutations upstream of the PI3K/Akt insulin signaling pathway were responsive to dapagliflozin, while those driven by mutations downstream of PI3K/Akt or in pathways with other driver mutations did not. These data support the development of precision medicine, insulin-lowering approaches to breast cancer in hyperinsulinemic patients, with or without diabetes, and position SGLT2 inhibitors as an attractive target to fill this niche.

## MATERIALS AND METHODS

### Study Design

4T1 (CRL-2539), Ac711 (CRL-3092), EMT6 (CRL-2755), EpH4 1424 (CRL-3071), M158 (CRL-3086), and M6 (CRL-3441) cells were purchased from ATCC and cultured in the manufacturer’s recommended media. These cells were injected into mice as described below at passage <10. In the cell division assays, 4T1 cells were cultured in the manufacturer’s recommended media containing 0.5% DMSO and supplemented with palmitate (0.5 mM), etomoxir (0.2 mM), and/or dapagliflozin (100 μM). Cells were counted by a blinded investigator using a LUNA-II automated cell counter after staining with Trypan blue 48 hours later.

All mouse studies were approved by the Yale University Animal Care and Use Committee. To determine the appropriate diabetes drug to lower both fasting and postprandial plasma insulin concentrations, female C57bl/6J mice were fed a high fat, high carbohydrate Western diet (Research Diets D12492; 60% calories from fat, 20% protein, 20% carbohydrate; 5.21 kcal/g in combination with 5% sucrose drinking water) for four weeks, after which two weeks of antihyperglycemic drug treatment were initiated. Mice were randomized to receive either vehicle (regular water and daily intraperitoneal injections of PBS), metformin (1 mg/ml in drinking water; approximate daily dose 200 mg/kg), dapagliflozin (0.0125 mg/ml, approximate daily dose 2.5 mg/kg), semaglutide (50 nmol/kg/day by subcutaneous injection), pioglitazone (10 mg/kg/day by intraperitoneal injection), or glipizide (0.5 mg/ml; approximate daily dose 100 mg/kg). After two weeks, blood glucose and plasma insulin were measured in the same mice after fasting for 4 and 16 hrs.

For the tumor studies, C57bl/6J, BALB/c, FVB, and MMTV-PyMT mice were purchased from Jackson Laboratories at 7 weeks of age. Upon arrival, a randomly selected subset was given a Western diet, while the remaining mice were given regular chow (ENVIGO-Teklad 2018; 18% calories from fat, 24% protein, 58% carbohydrate; 3.10 kcal/g) and regular water. Water intake was monitored twice weekly for each cage of five mice by weighing water bottles. Two weeks after arrival, mice were injected with tumor cells into the right mammary fat pad as shown in Supplementary Table 1.

All mice – whether they had spontaneous or orthotopic tumors – were monitored twice weekly until they developed palpable tumors, and daily thereafter. Chemotherapy (intraperitoneal paclitaxel, 15 mg/kg twice weekly) was initiated when tumors reached 300 mm^3^, measured using calipers. A subset of mice in both the chow and Western diet fed groups was randomized to receive dapagliflozin in drinking water beginning the day of chemotherapy induction, with the dapagliflozin concentration adjusted for the calculated daily water intake so that each mouse would receive approximately 2.5 mg/kg/day. A subset of otherwise untreated, tumor-bearing mice was subjected to acute dapagliflozin treatment: two doses of dapagliflozin (2.5 mg/kg per dose), each a day apart, by oral gavage. Terminal studies were performed 6 hr after the second dose.

In 4T1 and MMTV-PyMT tumor-bearing mice, dapagliflozin-treated mice were further randomized to be implanted with an Alzet pump containing insulin in artificial plasma (115 mM sodium chloride, 5.9 mM potassium chloride, 1.2 mM magnesium chloride hexahydrate, 1.2 mM sodium phosphate monobasic monohydrate, 1.2 mM sodium bisulfate, 2.5 mM calcium chloride dihydrate, 25 mM sodium bicarbonate, 4% bovine serum albumin) or vehicle (artificial plasma). The infusion rates were selected to match plasma insulin concentrations in 4 hr fasted dapagliflozin-treated mice to those measured in the diet-matched controls: 18 mU/kg/hr in chow fed mice, and 40 mU/kg/hr in Western diet fed mice. Mice with 4T1 and MMTV-PyMT tumors were monitored, and tumor size measured by a blinded investigator, twice weekly. Body weight was measured weekly. Food intake and activity were measured in Columbus Lab Animal Monitoring System metabolic cages and averaged over three days during week 3 of treatment. Thermal algesia was assessed by placing mice on a hot plate (IITC Life Sciences) at 52°C. Thermal latency was defined as the time elapsed before the mouse withdrew its back paw(s), jumped, or scaled the wall of the hot plate enclosure. Mice with Ac711, EMT6, Eph4 1424, and M158 tumors were monitored daily to determine survival. Failure to right was considered a surrogate marker for mortality.

### Flux Studies

One week prior to terminal isotope tracer studies, mice underwent surgery under isoflurane anesthesia to place a catheter in the right jugular vein. Mice were allowed a week of post-surgical recovery, during which they received daily intraperitoneal non-steroidal anti-inflammatory (Carprofen, 5 mg/kg per day) providing analgesic coverage for the first 72 hr after surgery. Terminal studies were performed following a 6 hr fast. Animals were placed in restrainers and their tails secured lightly using tape, allowing them some space to move around. After 2 hr to acclimate to the restrainer, a 1.0 mCi bolus of [2-^14^C] 2-deoxyglucose was administered through the catheter. Blood was obtained from the tail vein every 10 min following 2-deoxyglucose injection, and blood glucose concentrations were measured using a handheld glucometer. 60 min after radioisotope tracer injection, mice were euthanized using IV Euthasol (50 mg/kg). Tumors were freeze-clamped in tongs prechilled in liquid nitrogen and stored at −80°C pending further analysis.

Glucose uptake was measured as described previously *(50, 51)*. Briefly, [^14^C] concentrations were measured using the Hidex 300 SL scintillation counter in plasma (tracer, [^14^C] 2-deoxyglucose) every 10 min after radioisotope tracer injection, and in tumor ([^14^C] 2-deoxyglucose-6-phosphate, a trapping pool which cannot be further metabolized) 60 min after injection. The plasma [^14^C] concentrations were fitted to an exponential decay curve using Microsoft Excel, and the following equation was used for calculation of tumor glucose uptake:

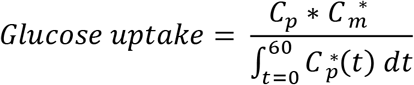

where C_p_ is the average plasma glucose concentration at the 30, 40, 50, and 60 min time points, C_m_* is the tumor [^14^C] (i.e., [^14^C] 2-deoxyglucose-6-phosphate) specific activity, and C_p_*(t), used in the integral, is the plasma [^14^C] 2-deoxyglucose concentration at time t.

### Biochemical Analysis

Blood glucose concentrations were measured using a handheld glucometer. Plasma insulin concentrations were measured by ELISA (Mercodia), and urine glucose concentrations using the YSI Glucose Analyzer. Plasma b-hydroxybutyrate (BOHB) concentrations were measured by gas chromatography/mass spectrometry (GC/MS), modifying a protocol we have previously described *(52)*. Plasma samples were spiked with an equal volume of internal standard [U-^13^C_4_] BOHB (1.0 mM), deprotonized with equal volumes of zinc sulfate followed by barium hydroxide, and centrifuged at 10,000 rpm for 10 min. The supernatant was transferred to a glass GC/MS vial and derivatized with 3 volumes of b-butanol 4N HCl, after which the samples were heated to 65°C for 60 min, evaporated under N_2_ gas, and resuspended in 75 mL of trifluoroacetic acid:methylene chloride (1:7). BOHB concentrations were measured by GC/MS run in chemical ionization mode, with peak areas for ^12^C BOHB compared to peak areas for ^13^C_4_ BOHB at a known concentration.

### Statistical Analysis

Survival analyses were performed using the Mantel-Cox logrank test, adjusting for multiple comparisons. Tumor size, fluxes, and biochemical parameters were compared by ANOVA with Tukey’s multiple comparisons test. The mean±S.E.M. are shown.

## Acknowledgments

The authors thank members of the Perry lab for helpful discussions, and members of the Yale Animal Resource Center for their care of the mice studied in these studies.

## Funding

Lion Heart Award (RJP)

National Institutes of Health grant T32-GM0007324 (supporting NDA)

China Scholarship Council (supporting XZ)

Yale Westchester Alumni Association (AAH)

Timothy Dwight Richter Fellowship (AAH)

## Author contributions

Conceptualization: NDA, RJP

Investigation: NDA, AN, WZ, AH, XZ

Funding acquisition: MBL, RJP

Project administration: RJP

Supervision: RJP

Writing – original draft: NDA, RJP

Writing – review & editing: NDA, AH, XZ, JRF, MBL, RJP

## Competing interests

RJP has previously received investigator-initiated research funding, for a project related to SGLT2 inhibitors but unrelated to cancer, from AstraZeneca.

## Data and materials availability

All data and materials used in the analysis are available to any researcher for purposes of reproducing or extending the analysis. Raw data are available without restrictions from the corresponding author upon request.

## Notes

https://www.proteinatlas.org/about/licence

