## Supplementary Tables and Figures for "A Precision Medicine Approach to Metabolic Therapy for Breast Cancer in Mice"

**Supplementary Materials**

| Cell Line | Mouse Strain | Cells injected |
|---|---|---|
| 4T1 | BALB/c | $1x10^4$ |
| Ac711 | C57bl/6J | $1x10^6$ |
| EMT6 | BALB/c | $1x10^6$ |
| Eph4 1424 | BALB/c | $5x10^5$ |
| M158 | C57bl/6J | $1x10^6$ |
| M6 | FVB | $2x10^5$ |

**Supplementary Table 1.** Details of the subcutaneous tumor cell injection protocols.

**Table S1.** Details of the subcutaneous tumor cell injection protocols.

| Cell Line | Strain | Source | Mutation(s) |
|---|---|---|---|
| 4T1 | BALB/c | ATCC | p53, Pik3cg |
| Ac711 | C57bl/6J | ATCC | HRAS |
| EMT6 | BALB/c | ATCC | PTEN |
| Eph4 1424 | BALB/c | ATCC | MEK1 |
| M158 | C57bl/6J | ATCC | MYC |
| M6 | FVB | ATCC | P53 |
| MMTV-PyMT | FVB | Jackson Labs | ERBB2, p53 |

**Supplementary Table 2.** Details of the models tested in this study.

**Table S2.** Details of the models tested in this study.

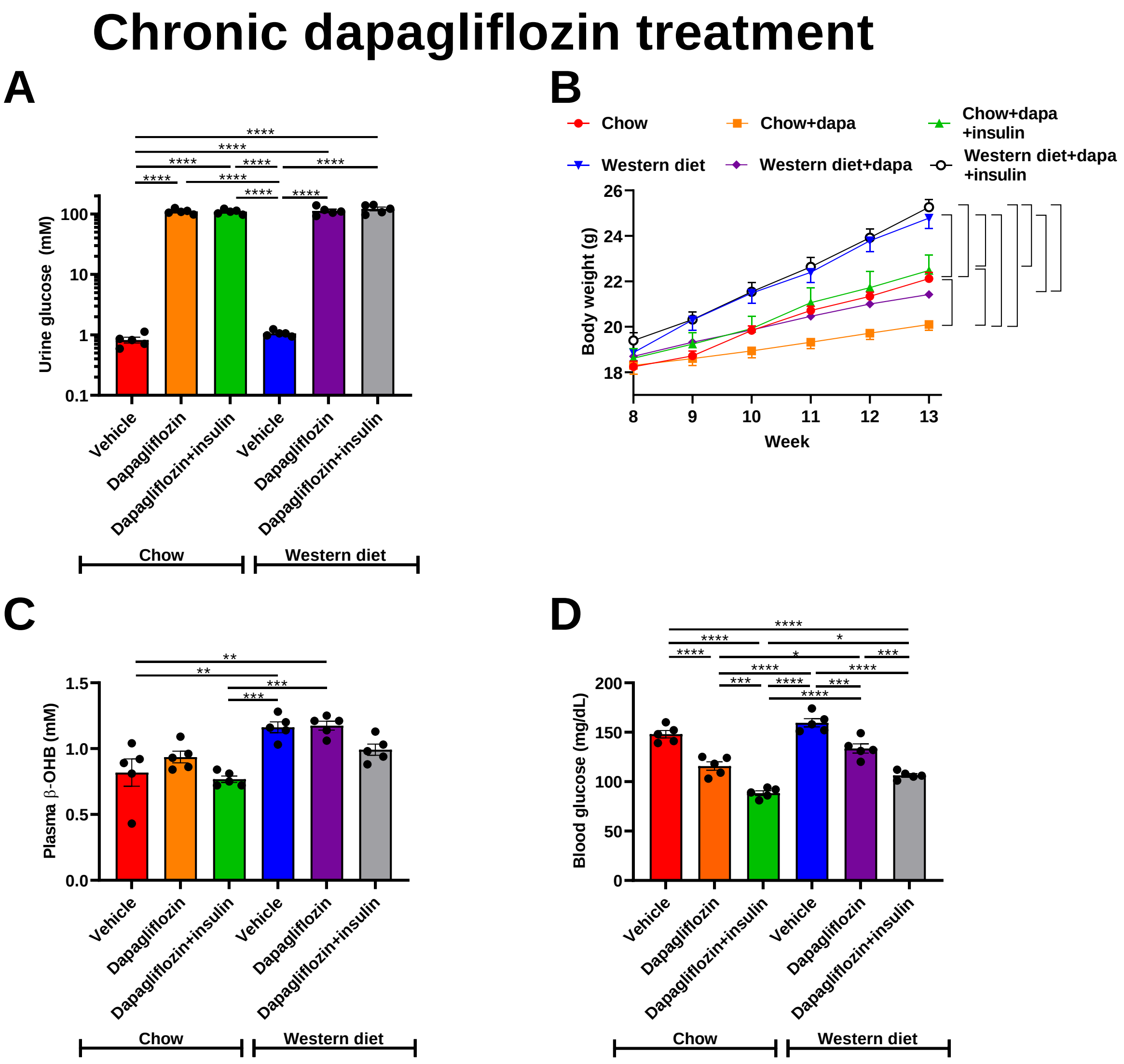


**Supplementary Figure 1. Dapagliflozin slows tumor growth in both lean and obese MMTV-PyMT mice in an insulin-dependent manner.** (A) Urine glucose. (B) Body weight. Brackets indicate statistically significant (*P*<0.05) comparisons. The mean±S.E.M. of n=5 per group is shown. (C) Plasma BOHB. (D) Blood glucose. In all panels, **P*<0.05, ***P*<0.01, ****P*<0.001, *****P*<0.0001 by ANOVA with Tukey's multiple comparisons test.

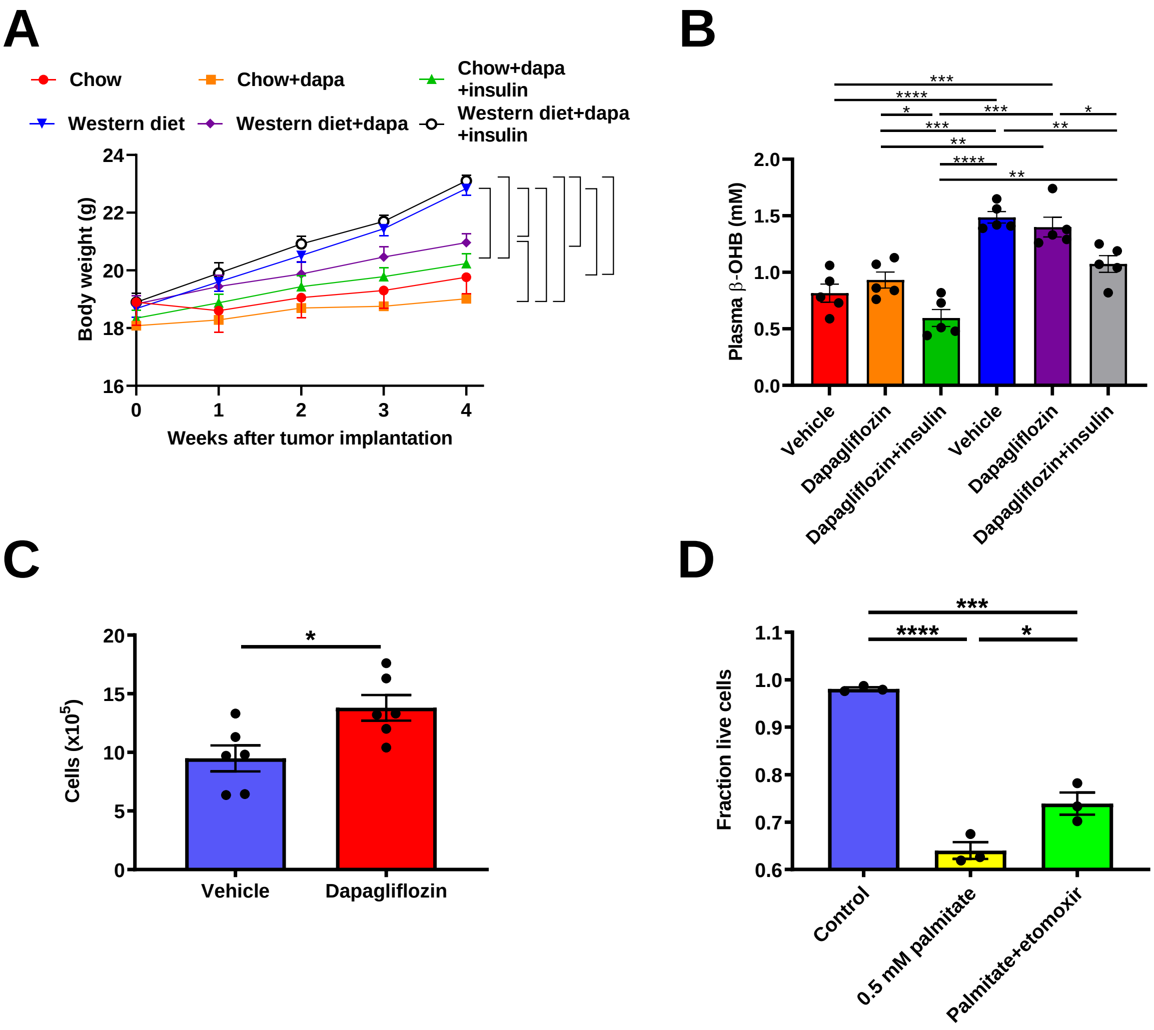


**Supplementary Figure 2. Dapagliflozin slows tumor growth in both lean and obese 4T1 tumor-bearing mice in an insulin-dependent manner.** (A) Body weight. Data are the mean±S.E.M. of n=5 per group. Brackets denote statistically significant (*P*<0.05) comparisons. (B) Plasma BOHB. (C) 4T1 cells cultured in vehicle (0.5% DMSO) or dapagliflozin (100 μM). **P*<0.05 by the 2-tailed unpaired Student's t-test (D) Fraction live 4T1 cells cultured in palmitate with or without etomoxir. In panels (A), (B), and (D), **P*<0.05, ***P*<0.01, ****P*<0.001, *****P*<0.0001 by ANOVA with Tukey's multiple comparisons test.
